# Precursors to Chunking Vanish as Working Memory Capacity Is Exceeded

**DOI:** 10.1101/515601

**Authors:** Regina Ershova, Eugen Tarnow

## Abstract

It is known that working memory capacity is limited. But what exactly happens if it is exceeded? Is the excess simply dropped or does it affect the items that are within the capacity of working memory? Here we show that as the capacity of working memory is exceeded, a rudimentary type of learning vanishes, namely, the probability of pairing items in the order of presentation.

Free recall of 500 Russian college students was measured using the Tarnow Unchunkable Test (Tarnow, 2014) consisting of sets of 3 and 4 double digit items. The average working memory capacity is exceeded with four items.

In the three item test, even though items were constructed to be unchunkable, there were asymmetric associations: recalling item N was more sensitive to whether item N-1 is recalled than the other way around. These asymmetric associations are presumably precursors of “chunking” and learning.

The asymmetric associations between items 1 and 2 and items 2 and 3 were similar. As the working memory capacity is exceeded in the four item test, the asymmetric association for the subject group halved from item 1 to item 2 (p=0.32) and disappeared completely from items 2 to 3 (large effect size: η^2^ =0.79, p=0.001) and there were no asymmetric associations from items 3 to 4.

## Introduction

Aristotle and the British Empiricists assumed that the human brain starts out as a tabula rasa (blank slate) and that, among the acquired content, contiguous stimuli create associations; this is referred to as the Law of Contiguity, (see Lachnit, 2003).

A measure in line with Hebbian learning is the sensitivity of the recall of one item to whether the previous item was recalled or not. This sensitivity is in part a contribution from whether the subject is attending to the items and from associations formed between the items. The former and some of the latter are symmetric with respect to item exchange (item N influences item N+1 the same way item N+1 influences item N); none of the former but some of the latter are asymmetric with respect to item exchange.

In this contribution we will examine how the limited working memory capacity (WMC; see Miller, 1956 and Cavanagh, 1972) impacts the symmetric and asymmetric associations between displayed items after single exposures to the item lists.

Previous work on inter-item associations in free recall has been focused on <u>static</u> associations, i.e. the probability of one item eliciting another as a free associate (see Deese, 1959 and citations thereof), and on fMRI studies of recognition probes of <u>dynamic</u> associations uncovering correlations between activity of different brain structures and current inter-item associations (see, for example, Piekerna et al, 2010),

## Method

We present data from a study of university students aged 17 to 24.

The Tarnow Unchunkable Test (TUT) used in this study is test of working memory capacity which uses lists of particular double-digit combinations which lack intra-item relationships (Tarnow, 2014). It does not contain any explicit WM operations. The TUT was given via the internet using client-based JAVAScript to eliminate any network delays. The instructions and the memory items were displayed in the middle of the screen. Items were displayed for two seconds without pause. The trials consisted of 3 or 4 items after which the subject was asked to enter each number remembered separately, press the keyboard enter button between each entry and repeat until all the numbers remembered had been entered. Pressing the enter button without any number was considered a "no entry". The next trial started immediately after the last entry or after a "no entry". There was no time limit for number entry. Each subject was given six three item trials and three four item trials in which the items are particular double-digit integers.

For each consecutive pair of items we calculate two associations:

> Forward association= probability that item N+1 is recalled given that item N is recalled - probability that item N+1 is recalled given item N is not recalled
>
> Backward association= probability that item N is recalled given that item N+1 is recalled - probability that item N is recalled given item N+1 is not recalled

We define

> Symmetric association=Average (Forward association, Backward association)

and

> Asymmetric association=Forward association - Backward association

Note that in our experiment the items are overwhelmingly recalled in order.

500 Russian undergraduate students of the State University of Humanities and Social Studies (121 (63%) females and 71 (37%) males, mean age was 18.8 years) participated in the study for extra credit. Each participant was tested individually in a quiet room. An experimenter was present throughout each session. One record was discarded - the student had only responded once out of a possible thirty times. This is an exploratory study.

## Results

The values of the symmetric and asymmetric associations for the 3-item and 4-item tests are displayed in Fig. 1, top row. While the average symmetric association is similar for both the 3-item and 4-item tests (ANOVA yields p=0.61 for item 1 to item 2 and p=0.76 for item 2 to item 3), the asymmetric association is much larger in the 3-item test than in the 4-item test: the asymmetric association is halved from item 1 to item 2 (p=0.32) and disappears completely from items 2 to 3 (large effect size: η^2^ =0.79, p=0.001) and from items 3 to 4. As expected, the asymmetric associations are positive, i.e. they go in the forward direction. The trial by trial symmetric and asymmetric associations for the 3-item and 4-item tests are displayed in Fig. 1), middle and bottom rows.

**Figure 1.**
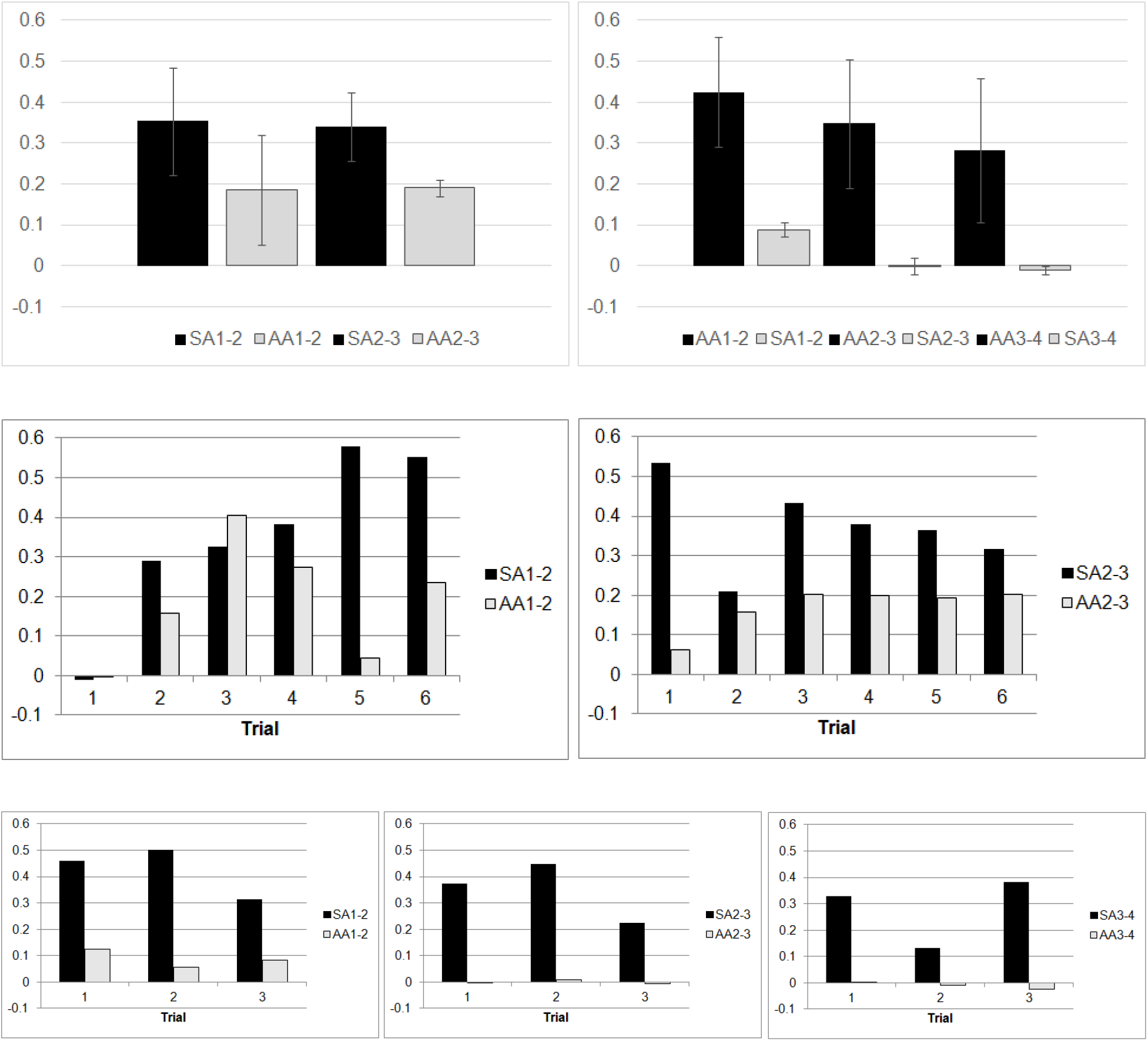
Top row: Symmetric association and asymmetric association for the 3-item (top left) and 4-item (top right) tests. SAM-N indicates the <u>s</u>ymmetric <u>a</u>ssociation between items M and N; AAM-N indicates the <u>a</u>symmetric <u>a</u>ssociation between items M and N. The error bars are the standard deviation of the results for the different trials. Middle & bottom rows: Symmetric and asymmetric association for the six 3- item trials for items 1 and 2 (middle left) and items 2 and 3 (middle right); for the three 4-item trials for items 1 and 2 (bottom left), items 2 and 3 (bottom center) and items 3 and 4 (bottom right).

If the asymmetric association disappears, does that mean the memory for item order disappears? In Fig. 2 is displayed the output order of subsequently displayed items. Display order is defined as +1 and reverse display order is −1. The graph shows that the vast majority of items are displayed in order, even for items 2-3 and 3-4 in the 4-item trials.

**Figure 2.**
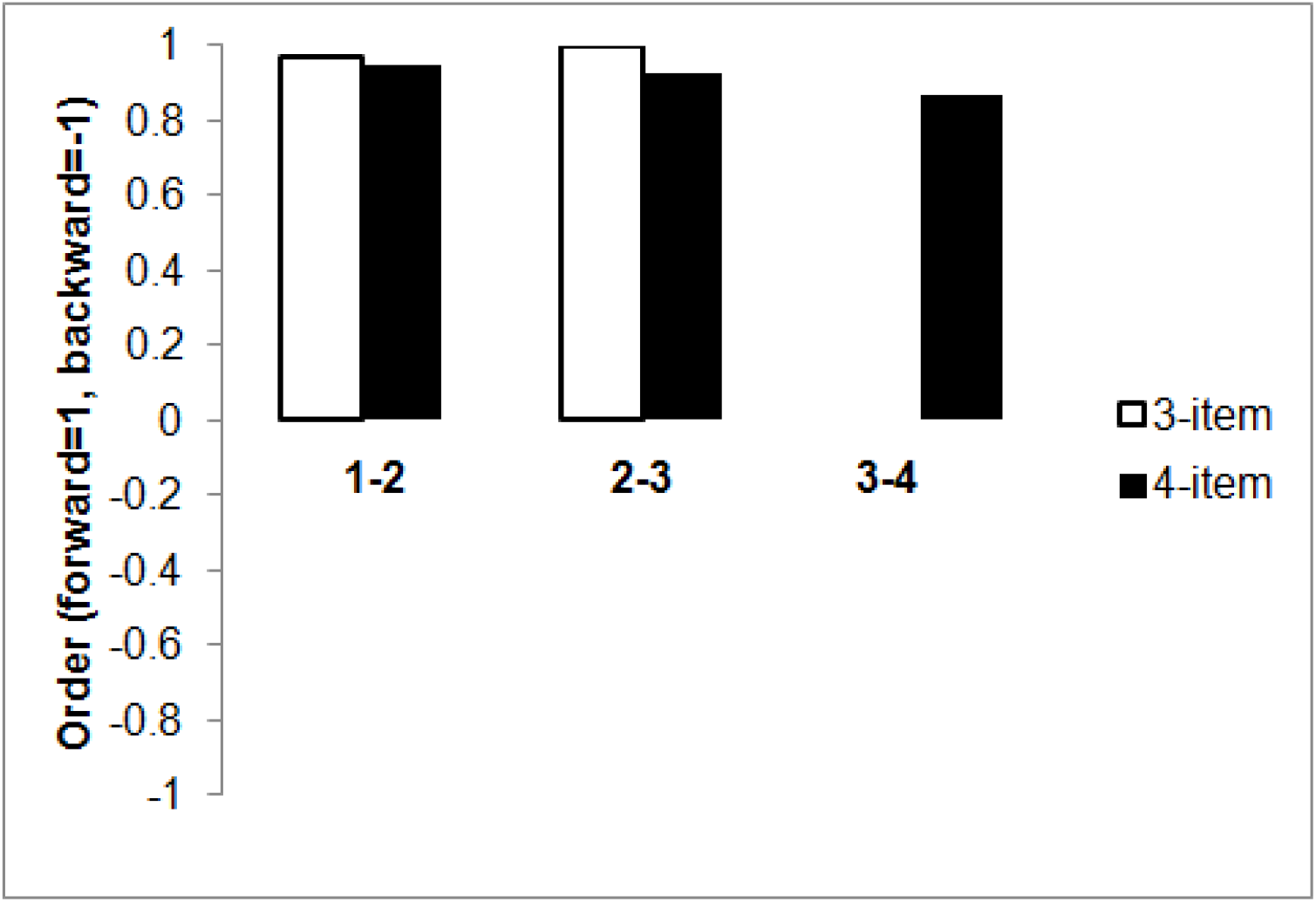
Recall order of subsequently displayed items.

## Discussion

We found before that WMC is limited to three TUT items (Ershova & Tarnow, 2016). Here we found that once the WMC is exceeded, the asymmetric associations between items disappear even while the symmetric associations remain. The effect size of our finding is very large with η^2^ =0.79. It lends credibility to the idea that information overload makes learning more difficult and stresses our message to textbook publishers - test the lengths of information lists.

The finding also provides yet another clue as to how working memory capacity is limited. We found earlier that the fourth item destroys the memory of the previous items (Ershova & Tarnow, 2018) - here we find that the extra item also removes the asymmetric associations.

The disappearance of asymmetric association curiously does not manifest itself in removing the item order in the output (Fig. 3). Others have noted that the memory for item order, as seen in an analysis of errors, is arguably not tied to a “chaining” algorithm (Henson, 1998); our result proves these findings in a very clear experiment.

This allows us to construct a tentative sensitivity chart for what happens when WMC is exceeded:

1. Most sensitive: Asymmetric associations between items disappear.
2. Sensitive: The number of recalled items cannot be increased and sometimes is decreased.
3. Least sensitive: Order memory stays similar to before WMC was exceeded.

